# A screening setup to streamline *in vitro* Engineered Living Material cultures with the host

**DOI:** 10.1101/2024.08.22.609167

**Authors:** Krupansh Desai, Shrikrishnan Sankaran, Aránzazu del Campo, Sara Trujillo

## Abstract

Engineered living materials (ELMs), which usually comprise bacteria, fungi, or animal cells entrapped in polymeric matrices, offer limitless possibilities in fields like drug delivery or biosensing. To determine the conditions that sustain ELM performance while ensuring ELM-host compatibility is essential before testing them *in vivo*. This is critical to reduce animal experimentation and can be achieved through *in vitro* investigations. Towards this goal, we designed a 96-well plate-based screening method to streamline ELM growth across culture conditions and determine their compatibility potential *in vitro*. We showed proliferation of three bacterial species encapsulated in hydrogels over time and screened six different cell culture media. We fabricated ELMs in bilayer and monolayer formats and tracked bacterial leakage. After screening, an appropriate medium was selected that sustained growth of an ELM, and it was used to study cytocompatibility *in vitro*. ELM cytotoxicity on murine fibroblasts and human monocytes was studied by adding ELM supernatants and measuring cell membrane integrity and live/dead staining, respectively, proving ELM cytocompatibility. Our work illustrates a simple setup to streamline the screening of compatible environmental conditions of ELMs with the host.

## 1. Introduction

Engineered Living Materials (ELMs) lie at the interface between life sciences and materials science [1]. In the biomedical field, these materials have been developed as wearables with biosensing capabilities, and as drug eluting devices with therapeutic and/or regenerative functions [1–3]. For example, probiotic bacteria embedded in a gastrointestinal device were developed as biosensors to detect inflammation in the gut [4]. Engineered *Bacillus subtilis* encapsulated in a poly (ethylene) glycol-poly (vinyl) alcohol (PEG-PVA) hydrogel were used as biosensors to detect isopropyl β-d-1-thiogalactopyranoside using a smartphone-based readout [5]. Engineered *E. coli* encapsulated in PDMS capsules could sense internal bleeding in the gastrointestinal tract of swine using heme detection [6]. In our recent work, *Corynebacterium glutamicum* engineered to produce hyaluronic acid and embedded in PVA hydrogels were used to form a self-replenishable lubricating layer on the surface of a contact lens [7]. Therapeutic living materials with controlled drug release were demonstrated with *ClearColi* engineered to produce deoxyviolacein upon blue light irradiation within Pluronic hydrogels [8, 9]. A similar approach was also used to release a pro-angiogenic protein and trigger differentiation in vascular endothelial cells controlled by light [10].

When it comes to application, the use of engineered living bacteria in ELMs raises concerns related to biosafety and compatibility with the host [1]. A few studies have investigated ELM-host interactions *in vitro*. Yanamandra *et al.* revealed that immune signaling pathways in healthy human peripheral blood mononuclear cells are not activated when co-cultured with ELMs containing *ClearColi* in Pluronic-based hydrogels [11]. Duraj-Thatte *et al.* showed *in vitro* compatibility of ELMs incorporating genetically modified probiotic *E. coli* that self-produced a mucoadhesive curli protein hydrogel. They tested the cytocompatibility of the ELMs towards Caco-2 cells by performing an MTT assay *in vitro*. The following *in vivo* experiments demonstrated that 10 days after ELM ingestion, gastrointestinal tract of mice showed no signs of strong immune response or epithelial damage in histopathology staining [12]. Liu et al. highlighted *in vivo* compatibility of living magnetic hydrogels incorporating genetically programmed *E. coli Nissle 1917* by histologically examining small intestine of mice treated with gel after oral ingestion. The staining confirmed no inflammatory response. Further, no drastic change in body weight and food-water uptake patterns of treated mice also underscored *in vivo* compatibility of these ELMs [13]. Lu *et al.* showed that mice treated with ELMs encapsulating *Lactococcus lactis* engineered to produce the vascular endothelial growth factor and lactic acid had negligible infiltration of inflammatory cells and massive collagen deposition in histological and Masson trichrome staining, respectively [14]. Praveschotinunt *et al.* demonstrated that their ELMs containing engineered *Escherichia coli Nissle 1917* to secrete curli fibers with trefoil factors were able to reduce mucosal damage and inflammation in the gastrointestinal tract. *In vivo* experiments showed that tissue cross-sections of mice treated with these ELMs had intact epithelial cell lining, lower edema and negligible infiltration of inflammatory cells at the site [15].

The living character of living therapeutic devices requires systematic testing for relevant assessment of their biocompatibility and safety in real application scenarios in contact with the body. Such experiments need to investigate the behavior of the engineered bacteria in the ELM and of the host cells as a function of tissue-dependent composition of the cellular milieu, as well as fluctuations in pH, oxygen or nutrient concentration. This requires a high number of co-culture experiments for which parallelized *in vitro* experimental formats and conditions need to be developed.

In this work, we report on important steps to develop predictive assays to test the biocompatibility and safety of ELMs *in vitro*. We describe a methodology to screen ELMs in 96-well plate format across different culture environments. We quantify bacterial proliferation inside the hydrogel, the leakage potential during culture time and the metabolic function of the bacteria inside the hydrogel. We assess the cytotoxicity potential using fibroblasts and monocytes. Our work will help to streamline the development of ELMs by optimizing early on the devices compatible with physiological conditions and thus, advancing ELM therapeutic investigations with the host.

## 2. Materials and Methods

The following reagents were used: LB powder (Lysogeny broth, Art. No. X964.2, Carl Roth GmbH, Germany), MRS powder (Man, Rogosa and Sharpe broth, Art. No. X925.2, Carl Roth GmbH, Germany), BHI powder (Brain heart Infusion, Art. No. X916.2, Carl Roth GmbH, Germany), GlutaMAX (Gibco GmbH, Cat. No. 35050-061, Germany), DMEM (Dulbecco’s Modified Eagle Medium) High Glucose (VWR GmbH, Cat. No. SIALD6429, Germany), Roswell Park Memorial Institute (RPMI) 1640 (RPMI (R), Gibco GmbH, Cat. No. 11835030), RPMI 1640 Medium ATCC modification (RPMI supplemented (R_supp_), Gibco GmbH, Cat. No. 1049101), Penicillin/ Streptomycin (Pen/Strep, Gibco GmbH, Cat. No. 15140-122, Germany), Trypsin-EDTA 1x (VWR GmbH, Cat. No. VWRL0154-0100, Germany), Fetal Bovine Serum (FBS) (PAN Biotech GmbH, Germany), vinyl sulfonated - poly (vinyl alcohol) (PVA-VS), LAP (Lithium phenyl-2,4,6-trimethylbenzoylphosphinate, Sigma Aldrich GmbH, Germany), Kanamycin (CAS No. 25389-94-0, Carl Roth GmbH, Germany), Erythromycin (Art. No. X4166.2, Carl Roth GmbH), glycerol (Art. No. 3783.1, Carl Roth GmbH, Germany), 3-(trimethoxysilyl) propyl acrylate (Art. No. 475149, Sigma Aldrich GmbH, Germany), OPI (Art. No. O5003, Sigma Aldrich GmbH, Germany), non-essential amino acids (NEAA, Cat. No.11140-050, Gibco GmbH, Germany), phosphate buffer saline (PBS, without Calcium and Magnesium, VWR GmbH, Germany), DAPI (4’,6-diamidino-2-phenylindole, dihydrochloride, Cas. No. 28718-90-3, Thermo Scientific GmbH, Germany), Tween 20 (Art. No. P1379, Sigma Aldrich GmbH, Germany). All these reagents were used without any modifications.

### 2.1 Bacteria inoculation

The bacterial strains used in this study were: *Escherichia coli* Nissle 1917 (EcN), *Lactiplantibacillus plantarum* WCFS1(LP), and *Corynebacterium glutamicum* (Cg) constitutively expressing the fluorescent protein, mCherry that is encoded in plasmids suitable for each strain. Plasmid maps for EcN, LP, and EcN are depicted in Figure **S1-3**. First, growth medium for each bacterial strain was prepared and sterilised by autoclaving. For EcN, LB was dissolved in MilliQ water (20 g/L) and supplemented with kanamycin (50 µg/mL). For LP, MRS broth was dissolved in MilliQ water (52 g/L) and supplemented with erythromycin (10 µg/mL). For Cg, BHI broth was dissolved in MilliQ water (37 g/L) and supplemented with kanamycin (50 µg/mL). Bacterial (either Cg, EcN or LP) inoculation was carried out by gently scratching the frozen bacterial culture in 20 wt.% glycerol (glycerol stocks stored at −80 °C) with a pipette tip. The pipette tip was then added to 5 mL of the growth medium. The bacteria were allowed to grow overnight, and optical density of the cultures were measured at 600 nm (OD600) using a spectrophotometer (Nanodrop 2000, Thermo Scientific GmbH, Germany).

### 2.2 Bacterial growth in suspension

Bacterial growth in different media was monitored for 15 to 20 hours. After inoculation bacterial cultures were diluted to an OD600 of 0.1 and centrifuged (3 min, 9000 rpm) to obtain a pellet. The bacterial pellets were resuspended in different media. EcN was cultivated in either LB broth or RPMI + 10% FBS (R-10%). LP was cultivated in MRS Broth, R-10%, RPMI supplemented + 10% FBS (R_supp_-10%) or RPMI supplemented + 20% FBS (R_supp_-20%). Cg was cultivated in either BHI, R-10%, R_supp_-10% or R_supp_-20%. Notably, Compared to RPMI® medium, RPMI ATCC Modified (R_supp_) medium contains HEPES, sodium pyruvate, high glucose content (4.5 g/L) and low sodium bicarbonate (1500 mg/mL).

To assess the possible effect of UV irradiation used for hydrogel polymerization on bacteria growth, 200 μL of bacterial culture or 200 μL of medium (negative control, blank) were transferred to a 96-well plate. The plate was irradiated with the conditions used for photo-crosslinking (3 min 15 s at 6 mW/cm^2^, 365 nm, UV lamp Bio link 365, Labor Technik GmbH, Germany). As controls, bacterial cultures without UV irradiation were used in the same plate. Bacterial growth was monitored for around 20 hours by measuring OD600 using a plate reader (TECAN Infinity, Germany). The plate reader was maintained at a different temperature depending on the bacterial strain (37°C for EcN and LP, 30°C for Cg). OD600 was read every 20 min at 4 different points in each well. Each measurement cycle started with shaking the plate for 18 min, followed by a rest period of 2 min and the OD600 measurement. Conditions were set in triplicates.

### 2.3 Bacterial preparation for encapsulation in hydrogels

After growing the bacteria overnight as described above, the cultures were diluted to OD 0.5, centrifuged, and resuspended in the appropriate media to mix with the polymer precursor solutions.

### 2.4 Hydrogel solutions preparation

PVA-VS (vinyl sulfone modified polyvinyl alcohol, 199-209 kDa, degree of VS-functionalization 0.95-1.34%) was synthesized as previously reported [7]. A 10 wt.% PVA-VS stock solution was prepared by stirring the polymer in MilliQ water at 1050 rpm at a temperature of 90-95°C during 3-4 h. PVA (130 kDa) was dissolved in a similar way at 10 wt.% concentration.

PVA hydrogels were prepared with either a bilayer architecture (i.e., inner layer encapsulating bacteria and outer protective layer preventing bacterial escape) or only inner layer to study effect of biocontainment. The inner layer was prepared by mixing PVA-VS at a final concentration of 4.75 wt.%, PVA at a final concentration of 0.25 wt.%, and LAP at a final concentration of 0.5 wt. % in the corresponding medium to be tested and the bacterial suspension at final OD600 0.05 in the same medium. The mixture was vortexed for 2 min at 3000 rpm. To prepare the outer protective layer for bilayer hydrogel, LAP at 0.5 wt.% final concentration was dissolved in 10 wt. % PVA-VS by sonicating for 15 min at room temperature and shaked in an Eppendorf ThermoMixer (40°C, 1200 rpm) for 15 min. The volumes for each condition tested are specified in Tables S1, S2, and S3. Hydrogels are named X-PVA, X being the bacterial strain used in the inner layer (EcN, LP or Cg). Controls gels without bacteria (OD 0) were prepared in both bilayer and inner layer architecture.

### 2.5 Hydrogel fabrication in µ-plate 96 well 3D

Bilayer hydrogels were prepared in µ-plate 96Well plates (Ibidi, Germany). The 96 wells had a diameter of 5 mm diameter (ca. 70 µL volume) and an internal microwell with 4 mm diameter and 0.8 mm height (ca. 10 µL volume). Prior to hydrogel preparation, the surface of the wells was functionalized with acrylate groups to covalently attach the PVA-VS hydrogel to the surface of the well. For this purpose, 70 µL of absolute ethanol was first added to the wells and the plate was kept on a shaker for 5 min at 300 RPM. After removing the ethanol, 70 µL of a 0.5% solution of 3-(trimethoxysilyl) propyl acrylate in ethanol were added to the wells and the plate was kept overnight on a shaker at 300 RPM. On next day, the plate was left under the fume-hood to let the remaining ethanol evaporate. Plates were UV-sterilized for 30 min in a safety laminar flow hood before use. 10 µL of the hydrogel precursor mixture were added to the inner well to obtain the inner layer and polymerized by UV irradiation (365 nm, 6 mW/cm^2^) for 45 s using a UV lamp (Bio link 365, Labor Technik GmbH, Germany). Afterwards, 20 µL were added to the well to form the outer protective layer and photo-crosslinked (365 nm, 6 mW/cm^2^) for 2 min and 30 s. After polymerization, 50 µL of medium was added. Plates were incubated at 37°C and 5% CO_2_ unless otherwise noticed (Binder, Germany).

### 2.6 Quantification of bacterial proliferation in the hydrogels

To quantify bacterial proliferation in the hydrogel, alamarBlue assay (Invitrogen, Germany) was used at different timepoints following manufacturer’s instructions. Briefly, medium was exchanged to medium with alamarBlue reagent at different timepoints. 10% alamarBlue reagent was added to the specific medium to test and hydrogels were incubated at 37°C (or 30°C when noted), 5% CO_2_ for 4 h with the reagent. After incubation, 25 µL of the medium was transferred to black 384-well plate and fluorescence was measured using a plate reader (Tecan Biotek, Ex/Em 530/570 nm and 580/610 nm). Samples were measured in triplicates.

### 2.7 Determination of bacterial leakage from the hydrogels

To test bacteria leakage from the hydrogels after culture, 2 μL of the cultured supernatant were transferred to a 96-well plate and 200 uL of bacterial growth media (MRS Broth supplemented with erythromycin for LP, LB broth supplemented with kanamycin for EcN and BHI medium supplemented with kanamycin for Cg). Positive controls with inoculated bacteria and negative controls with bacterial growth medium were used. These cultures were incubated overnight and the OD600 was measured at the beginning (0 h) and after 24 h or 48 h using a plate reader set at 37°C (for EcN) or 30°C (for Cg).

### 2.8 Fluorescence microscopy

All three bacterial strains expressed mCherry. For live imaging, a Zeiss Celldiscoverer 7 microscope installed with LSM 900 and Airyscan 2 (Zeiss, Germany) was used. Images were taken at 20x / 0.95 NA (ZEISS Plan-Apochromat) objective with optovar 1x or 0.5x Tubelens lens. While imaging, the temperature was set at 37°C. mCherry fluorophore was visualized with a Ex/Em of 587/565-700 nm and Z-stacks of 20 µm were captured at least 50 µm above the bottom of the plate. At least 3 stacks per replicate were taken and 3 replicates per condition were used.

Bacterial colony areas were calculated from stacks of 20 um at 10X (16-20 images in total). Stacks were projected onto the same plane using the *Z-projection* tool on Fiji (ImageJ v1.54f) and applying the *Max. Intensity* projection. From the projections, masks were obtained using the *threshold* tool (Otsu method, automatic detection) and then individual colonies were selected manually using the *wand* tool. For all timepoints >200 colonies were quantified.

For EcN samples, brightfield images were used at 10X (19 images per timepoint in total). To quantify the colony shape descriptors, masks were obtained first. The flatten calibration tool in Fiji was applied first. Then, the *auto-crop* tool was applied. After that, each image was converted to 8-bit and the *auto-threshold* was applied (Otsu method). Then, the tool *fill holes* was used and colonies on focus were selected with the *magic wand* tool and measured. For each timepoint measured >200 colonies were quantified.

### 2.9 Cell culture

Fibroblasts (NIH-3T3 cell line, ATCC, mouse, passages 22-27) were used for cytotoxicity assays. Cells were grown in DMEM high glucose and supplemented with 10% heat inactivated FBS (D-10%), 1% GlutaMax, and 1% Penicillin/Streptomycin. Cell culture was exchanged every other day and cells were passaged (trypsin-EDTA, 3 min at 37 °C, 5% CO_2_) when cell confluency reached around 70%. Fibroblasts were seeded at a final cell density of 16000 cells/cm^2^ in 96-well plates using growth medium (D-10% + 1% GlutaMax +1% Pen/Strep). After 24 h, the culture medium was replaced by supernatants from the bacterial gels (Cg-PVA), which were diluted in R_supp_-20% (1:1, V:V). Fibroblasts were incubated with supernatants for 24 h and then, lactate dehydrogenase (LDH) assay and alamarBlue assay were performed.

Monocytes (Mono-mac-6 cell line, ATCC, human, passages between 12-15) were used for cytotoxicity assays. Cells were grown in RPMI medium supplemented with 10% heat inactivated FBS, 1% GlutaMax, 1% NEAA, 1% OPI and 1% Pen/Strep. Cell culture was changed every other day and cells were diluted to 300,000 cells/mL every 3-4 days (counted as a passage).

Monocytes were seeded at a final cell density of 150,000 cells/mL in 96-well (Greiner Bio-one, Germany) plates using growth medium. After 24 h, the culture medium was replaced by supernatants from the bacterial gels diluted in R_supp_-20% (1:1, V:V). Monocytes were incubated with supernatants for 24 h and then, live/dead staining was performed.

### 2.10 Cytotoxicity assays

To perform the cytotoxicity assays, 50 μL of supernatants from Cg-PVA hydrogels incubated in R_supp_-20% and collected on day 1 and day 2 were used and mixed with 50 μL of R_supp_-20% medium.

Lactate dehydrogenase (LDH) assay (CytoTox 96^®^ Non-Radioactive, Promega, Germany) was performed to measure cell death following manufacturer’s instructions. Briefly, 30 μL of supernatants were transferred to a 96-well plate and mixed with 30 μL of the CytoTox 96® Reagent. The plate was incubated at room temperature (300 RPM, protected from light) for 30 min. After that, the stop solution reagent (30 μL) was added and absorbance at 490 nm was measured using a plate reader. Blanks of each supernatant were used, and lysed cells with 5 μL of 37.5% Triton X-100 were used as positive control. 5 μL of PBS was added to samples to keep the same final volume in all the wells.

AlamarBlue assay was performed to determine the number of metabolically active cells. Briefly, 10% alamarBlue reagent was added to cell culture medium. Cells were incubated with the medium supplemented with alamarBlue reagent for 1.5 h at 37°C (5% CO_2_). Then, 50 μL of supernatant were transferred to a black bottom 96-well plate. Fluorescence intensities were measured in a plate reader (Ex/Em 570/600 nm). Blanks with only supernatants and positive controls of cells grown in growth medium were included.

### 2.11 Immunostaining

After performing LDH and alamarBlue assay, fibroblasts were washed twice with PBS and fixed using 4% PFA (Paraformaldehyde solution) for 30 min at room temperature. After fixation, samples were washed twice with PBS to remove the PFA left. Cells were permeabilized using 0.1% of triton X-100 in PBS for 5 min at room temperature. Then, cells were washed twice with PBS and blocking buffer (1% Bovine serum albumin in PBS) was added for 30 min at room temperature. After removing blocking buffer, phalloidin (AlexaFluor 488-Phalloidin, 1:400 dilution, Proteintech, Germany) and DAPI (1:1000 dilution, Invitrogen, Germany) were added and incubated at room temperature for 1 h. Finally, cells were washed five times with wash buffer (0.5% Tween 20 in PBS) and cells were incubated in PBS while imaging. Zeiss Celldiscoverer 7 microscope installed with LSM 900 and Airyscan 2 (Zeiss, Germany) was used to capture fluorescence images at 20x / 0.95 NA (ZEISS Plan-Apochromat) objective with optovar 0.5x Tubelens lens.

### 2.12 Live/Dead staining

Monocytes were stained for viability determination after adding the bacterial gels supernatant overnight. Cells were washed twice with PBS. The live (fluorescein diacetate, 5 mg/mL, Invitrogen, Germany) and dead (propidium iodide, 2 mg/mL, Invitrogen, Germany) staining were added to PBS to make final concentration of 40 μg/mL and 30 μg/mL, respectively. The samples were incubated at room temperature for 15 min. Then, staining was removed, and the samples were washed twice with PBS before imaging. Finally, images were taken immediately using an epifluorescence microscope (Keyence). A minimum of 10 images at 10x magnification were obtained. Quantification of % of live cells was carried out using the 10x magnification images. The % of live cells was calculated as: [number of live cells/total cells] *100.

### 2.13 Statistical analysis

All experiments were performed in triplicates. Statistical analysis was carried out using GraphPad Prism v9 software. All graphs represent the mean ± standard deviation (SD) unless otherwise noted. The goodness of fit of the data sets was assessed by Normality Shapiro-Wilk test. For comparisons of three groups, normal distributed populations were analyzed via analysis of variance (ANOVA) test followed by a Tukey’s *post hoc* test to correct for multiple comparisons. For comparisons of two groups, normal distributed populations were analyzed via parametric t-test. For populations of two groups that were not normally distributed, a non-parametric (Mann-Whitney) t-test was performed. For comparisons of two groups at different timepoints, populations were analyzed via analysis of variance (two-way ANOVA). Two-way ANOVA was performed by matching timepoint as a factor. A full model was fitted (time (row) effect, condition (column) effect and time/condition (row/column) effect). An equal variability of differences was not assumed and therefore the Geisser-Greenhouse correction was used. Multiple comparisons were performed two ways, first within each condition through time and then comparing conditions per timepoint (column (condition) and row (time) effects). Differences among groups are indicated as follows: *p*-values <0.05 (*), *p*-values <0.01 (**), *p*-values <0.005 (***), *p*-values <0.001 (****), and differences among groups not statistically significant (ns).

## 3. Results

We investigated ELMs containing three different bacterial strains: two gram-positive strains (*Corynebacterium glutamicum* (Cg) and *Lactiplantibacillus plantarum* WCFS1 (LP)) and one gram-negative (*Escherichia coli* Nissle 1917 (EcN)). We selected LP and EcN because they are generally recognized as safe (GRAS) in the food industry [18, 19], they are natural commensals in humans [20, 21], and these probiotic bacteria are highly used in ELMs [11, 12, 14] and as living biotherapeutic products [22, 23]. We selected Cg due to our previous work on a living contact lens prototype [7], where the species *Corynebacterium* represents approximately 80% of the total eye microbiome [24, 25]. For imaging purposes, we used engineered variants of these bacteria strains that expressed mCherry.

We performed initial cytotoxicity assays on murine fibroblasts in LB, BHI or MRS, which are the broths used to culture the selected bacteria strains. The addition of LB, BHI or MRS broths for 24 h to the fibroblasts in culture led to an 8-, 6- and 3-fold decrease in cell viability as measured by alamarBlue assay compared to, for example, RPMI medium supplemented with 10% FBS (R-10%) (**Figure 1a**). Microscopic observations of cell morphology (**Figure S4**) showed a progressive rounding and detachment of the fibroblasts from the well plate when in contact with either LB, MRS or BHI, whereas they presented a spread morphology when cultured in R-10%, RPMI ATCC modified and supplemented with 10% FBS (R_supp_-10%) and DMEM supplemented with 10% FBS (D-10%).

**Figure 1.**
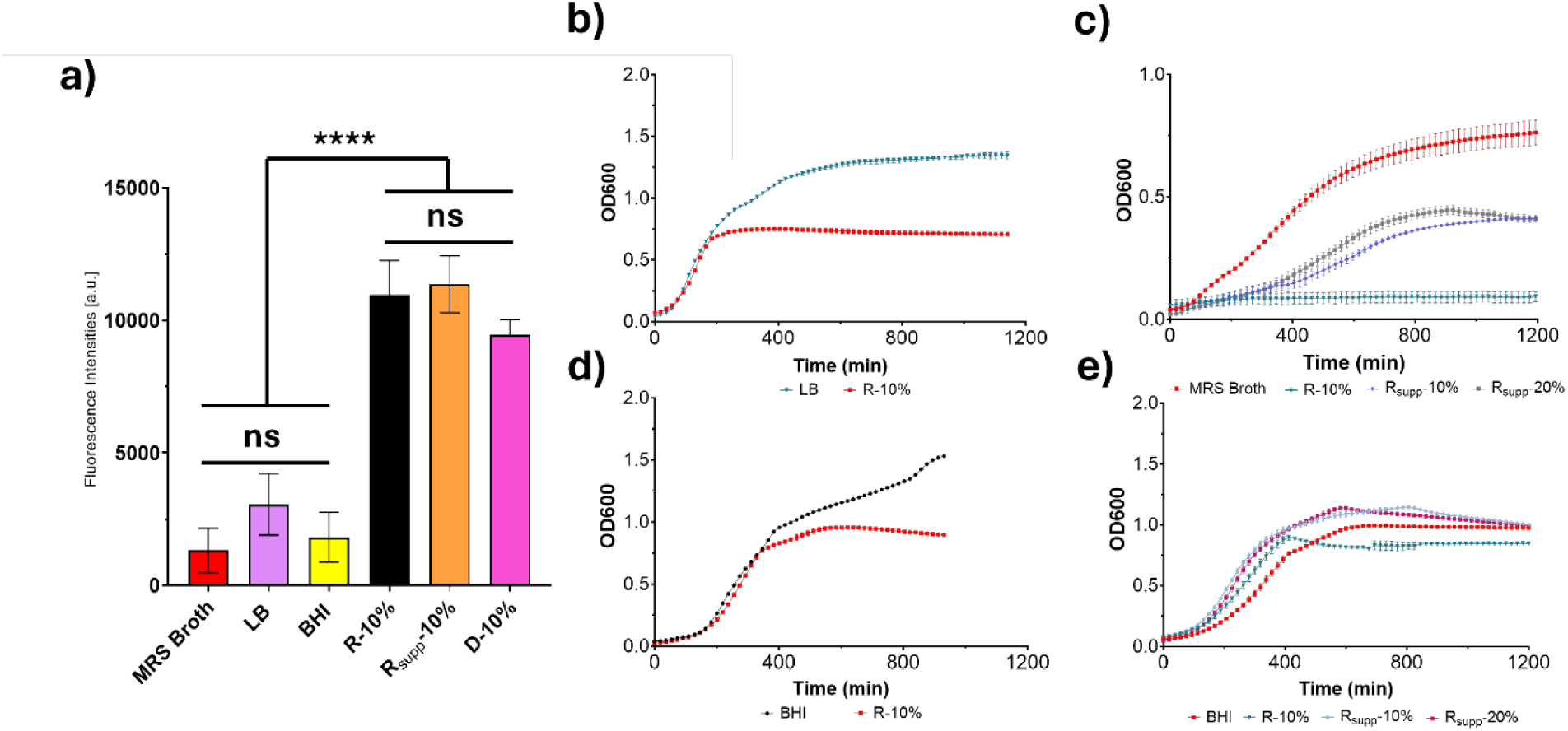
Mammalian cell culture conditions affect bacterial growth. **(a)** Viability of fibroblasts in contact with bacterial culture media (MRS Broth, LB, and BHI). R-10%, R_supp_-10% and D-10% were used as controls; **(b)** Growth curve of EcN in suspension at 37°C in LB broth (blue line) and RPMI-10% FBS (R-10%, red line **(c)** Growth curves of LP in suspension at 37°C in MRS broth (red line), R-10% (teal line), R_supp_-10% (grey line), R_supp_-20% (purple line) **(d)** Growth curve of Cg in suspension at 30°C in BHI broth (black line), R-10% (red line) **(e)** Growth curve of Cg in suspension at 37°C in BHI broth (red line), R-10% (teal line), R_supp_-10% (light blue line), R_supp_-20% (purple line). For all plots data is represented as mean ± SD and n= 3 biological replicates.

On the other hand, the growth of the bacterial strains (*Escherichia coli* Nissle (EcN), *Lactiplantibacillus* (LP) and *Corynebacterium glutamicum* (Cg)) in suspension in RPMI or DMEM supplemented with 10% FBS (R-10%, Rsupp-10% or D-10%) showed notable differences between bacterial strain growth and amongst cell culture media assayed (**Figure 1b-e**).

EcN grown in R-10% (**Figure 1b**) presented a similar lag phase and slightly lower average growth rate (0.104 hr^-1^) in the exponential phase compared to LB broth (0.124 hr^-1^) but the stationary phase in R-10% was achieved at a lower OD600. LP strain did not grow in R-10% (**Figure 1c**) and, therefore, we investigated growth in RPMI supplemented with high glucose (4.5 g/L), sodium pyruvate, and HEPES with either 10 or 20% FBS (R_supp_-10%, or R_supp_-20%). LP grew in both R_supp_-10% and R_supp_-20%. A longer lag phase compared to MRS broth was observed for both media and the exponential phase was achieved at a lower rate in both cases, R_supp_-10% being the slowest (average growth rate of 0.015 hr^-1^). The stationary phase reached by LP grown in R_supp_-10% and R_supp_-20% were at a lower bacterial density (OD600 0.41 and 0.37, respectively) compared to MRS broth (0.67).

We also screened different culture conditions in suspension with Cg (**Figure 1d, e**). Normally, this bacterium grows at 30°C, therefore, we started comparing the grow of Cg at 30°C in their optimal medium (BHI broth) and R-10%, which supported EcN growth but no LP (**Figure 1d**). At 30°C, the lag phase observed for BHI broth and R-10% was similar and the growth rate in exponential phase was similar (0.13 hr^-1^) for both conditions. Differences arose in the stationary phase. On average, Cg growing in R-10% reached stationary phase sooner compared to BHI broth, resulting on an overall lower final OD600 of 0.65. In BHI broth, Cg continued growing at a slower rate (average growth rate of 0.06 hr^-1^) compared to the exponential phase and by the end of the experiments Cg had not reached stationary phase. We also tracked the growth of Cg in suspension at 37°C. At 37°C, all media screened presented similar growth curves, with a similar lag phase. The exponential phase started approximately at the same time for all the media assayed with slight differences in the growth rate. For example, R_supp_-10% and R_supp_-20% presented a faster exponential growth (average growth rate of 0.127 hr^-1^ and 0.131 hr^-1^) compared to BHI (average growth rate of 0.087 hr^-^1) broth. Stationary phase was reached for all conditions tested, being R-10% the condition that obtained the lowest number of bacteria (OD600 of 0.89) at the end of the experiment.

The development of a more systematic screening method of the behavior of ELMs in different environments was motivated by i) the differences observed in bacterial growth in suspension in different media and ii) the viability decay observed in fibroblasts cultured in LB, MRS and BHI broths. This method aims to identify suitable conditions for co-culturing mammalian cells and bacteria-based ELMs.

We selected PVA-VS/PVA mixtures as hydrogel matrix for our ELMs since these have been successfully used by us and others in previous reports [7, 14, 16, 17]. Bilayer hydrogels were prepared in 96well-plates with a well-in-well architecture (**Figure 2a**). The bilayer hydrogels had an inner layer containing the bacteria and composed of 4.75 wt.% PVA-VS and 0.25 wt.% PVA (before swelling) and bacteria at OD0.05. The outer layer, with a protective function, did not contain bacteria and was composed of 10 wt.% PVA-VS (before swelling) (**Figure 2b**). These PVA/PVA-VS concentrations were selected considering our results in previous work with PVA-VS/PVA hydrogels for ELMs [7].

**Figure 2.**
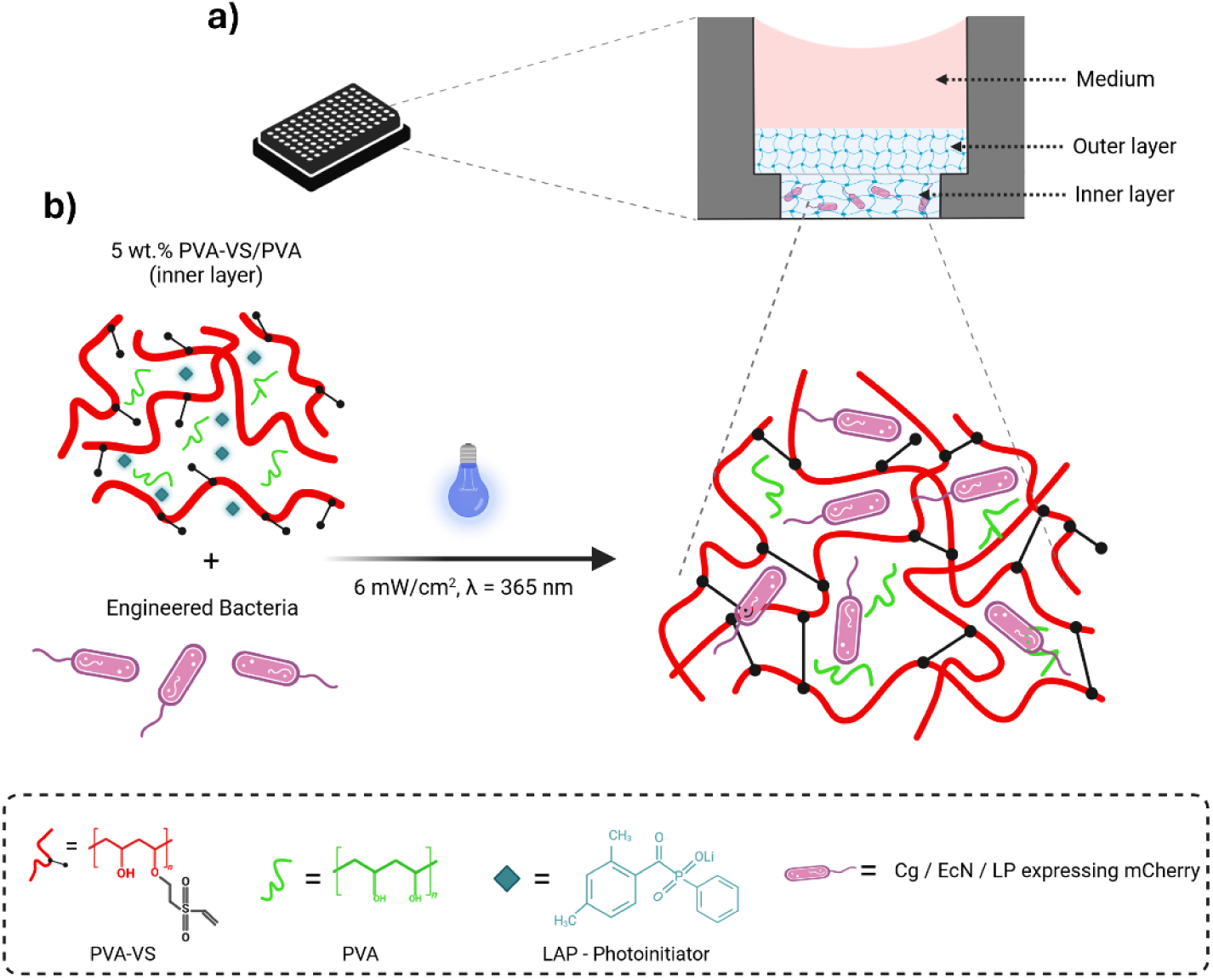
Composition of the bilayer hydrogel model inside the microwells. (**a**) PVA-VS/PVA bilayer ELMs fabricated in µ-plate 96 Well. (**b**) Photo crosslinking of bacterial PVA-VS/PVA hydrogels (inner layer) using UV radiation.

Prior to encapsulation in the hydrogels, we studied the effect of UV irradiation used to polymerize the hydrogels on bacterial viability in suspension cultures. No differences in bacterial growth were observed between LB and LB_UV irradiated, or R-10% and R-10%_UV irradiated (**Figure 1b** and **S5a**).

We studied viability and growth of EcN within the PVA hydrogels (EcN-PVA gels) in R-10% medium (**Figure 3**). We used alamarBlue assay to investigate EcN proliferation, as a measure of their metabolic activity. As shown in **Figure 3a**, bacteria grew inside the hydrogels from day 0 to day 1. We also tracked their growth via imaging as independent measure of bacterial proliferation. **Figure 3b** shows whole well images of the bacterial hydrogels with increasing culture time, where growing colonies can be observed. From these images, we quantified colony size and EcN formed colonies on day 1 that were bigger in size on day 2 (**Figure 3c**). EcN-PVA bilayer gels cultured in R-10% presented bigger and more sparse colonies compared to the LB broth control (**Figure S6**). As a measure of biocontainment, we tracked the occurrence of bacterial leakage from these ELMs by taking a small volume of the supernatant and using it to inoculate a suspension culture. Of all the EcN-PVA bilayer gels fabricated, 5% of the gels leaked on day 2 (**Figure 3d**).

**Figure 3.**
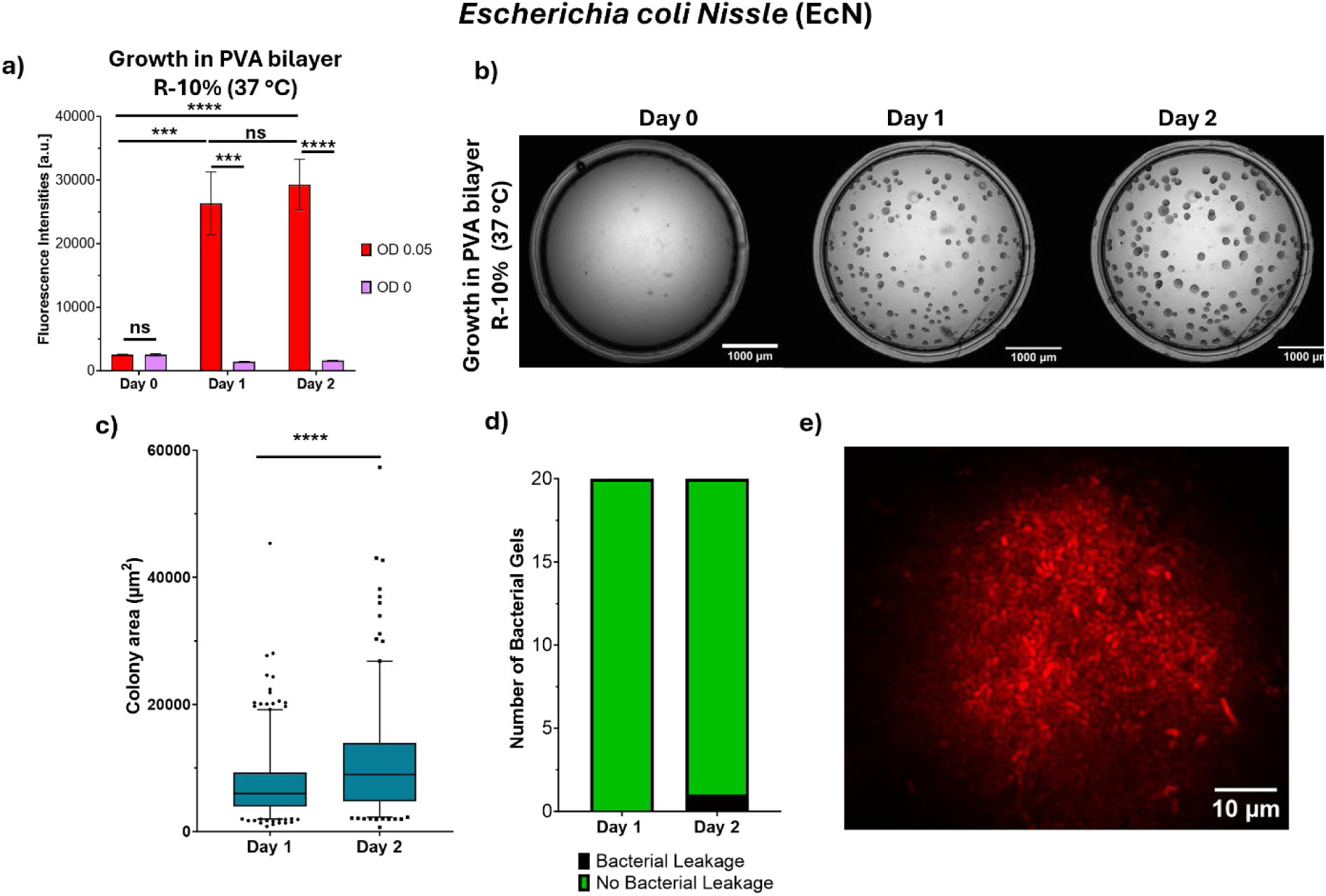
EcN encapsulated in PVA bilayer hydrogels grow in R-10% medium at 37°C. Bacterial hydrogels were fabricated by encapsulating a final OD600 of 0.05 and an empty hydrogel control (OD 0) was used. **(a)** Proliferation (manual gain 65, alamarBlue assay) of EcN encapsulated in PVA hydrogels (EcN-PVA gels) and incubated in R-10% at 37 °C (n ≥ 3 biological replicates, mean ± SD). **(b)** Bright field images of the whole wells containing EcN-PVA gels (scale bar: 1000 µm). **(c)** Colony area quantification (μm^2^) from EcN-PVA (The line within the box signifies the median value, and the whiskers denote the range between the 5^th^ and 95^th^ percentiles, with outliers depicted as dots positioned above or below the whiskers) **(d)** Quantification of leakage from EcN-PVA gels **(e)** Representative high magnification image of a single colony (Scale bar: 10 µm). Differences among groups are indicated as follows: p-values <0.05 (*), p-values <0.01 (**), p-values <0.005 (***), p-values <0.001 (****), ns = not significant.

To compare the functionality of the bacteria inside the different media, we tested protein production by imaging the fluorescence coming from the expression of mCherry (**Figure 3e**). High magnification images of the colonies showed homogeneous expression of mCherry within individual cells across the area of the colonies imaged.

Next, we encapsulated a Lactobacillus strain (LP) (**Figure 4**) and screened different cell culture media. Prior to encapsulation, we studied the effect of UV irradiation on LP growth in suspension (**Figure S5)**. UV irradiation for LP culture in MRS broth did not change its growth profile over time. Notably, LP did not grow in R-10%, irrespective of UV irradiation. UV treatment did not affect LP growth in R_supp_-10% or R_supp_-20%.

**Figure 4.**
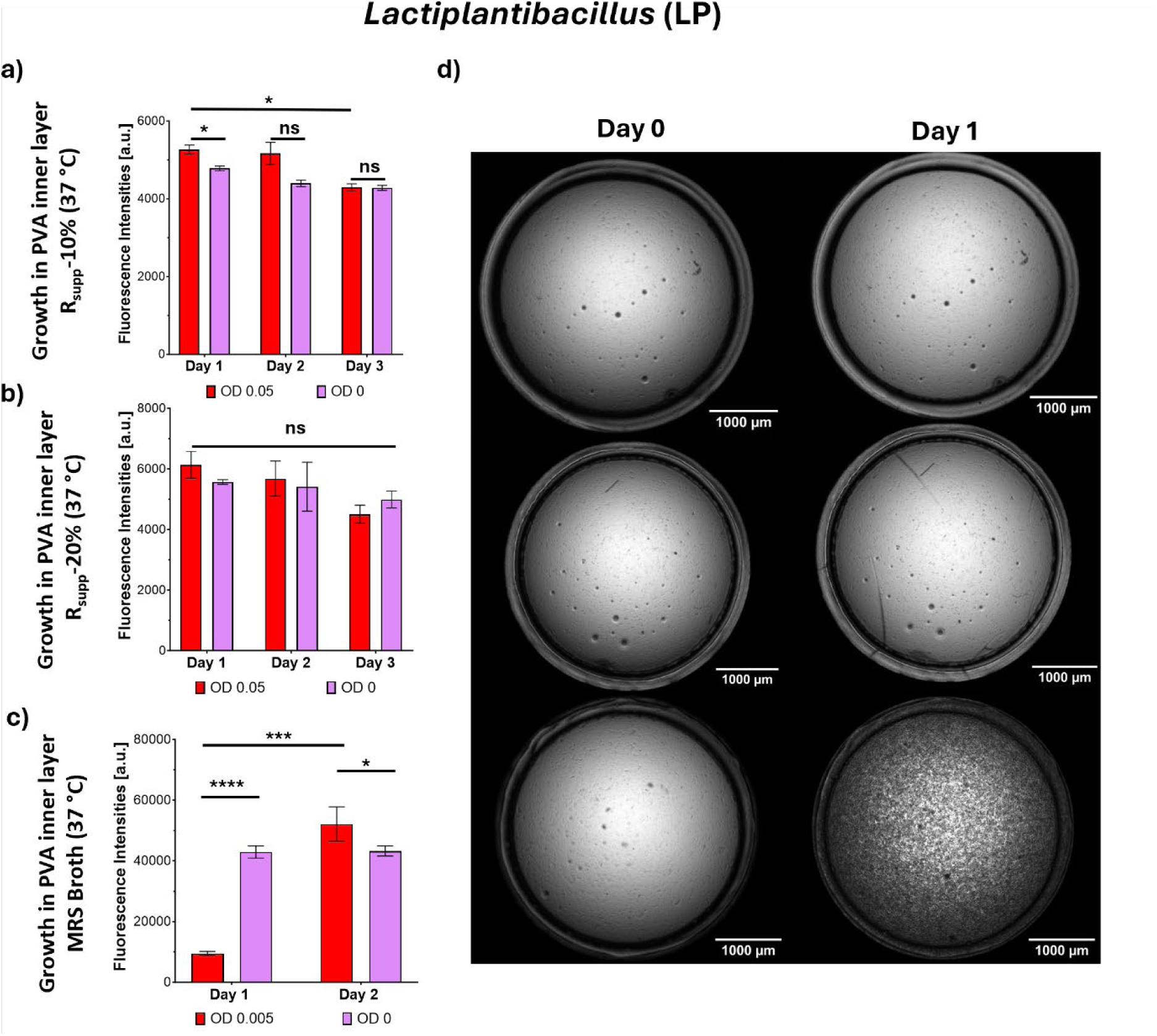
LP encapsulated in PVA inner layer hydrogels only grow in MRS broth at 37°C. Proliferation (manual gain 80, alamarBlue assay) of LP-PVA inner layer gels in **(a)** R_supp_-10% **(b)** R_supp_-20% and **(c)** MRS Broth at 37°C. (n= 3 biological replicates, mean ± SD) (d) Bright field images of inner layer gels encapsulating LP mCherry in R_supp_-10, R_supp_-20%, and MRS Broth on day 0 and day 1, respectively. Scale bars: 1000 µm. Differences among groups are indicated as follows: p-values <0.05 (*), p-values <0.01 (**), p-values <0.005 (***), p-values <0.001 (****), ns = not significant.

Then, we encapsulated LP in PVA bilayer gels and tracked their growth in R_supp_-10% and MRS broth, which allowed the growth of LP in suspension (**Figure 1c, Figure S7**). The proliferation of LP in LP-PVA bilayer gels was studied via alamarBlue assay (**Figure S7a**). No growth was observed in Rsupp-10% for up to 2 weeks. LP-PVA bilayer gels incubated in MRS broth (**Figure S7b**) presented high fluorescence values on day 1 but a steep decrease in fluorescence by day 2, measured by alamarBlue. The observations from brightfield or fluorescence images showed a different outcome (**Figure S7c**): here colonies were observed by day 1. Colony areas increased from day 1 to day 2 (**Figure S7d**) and LP were seen expressing mCherry homogenously across the colonies (**Figure S7e**). It is noteworthy that colony size for this strain was approximately 50-fold smaller compared to EcN grown in R-10%.

To investigate whether the presence of the outer hydrogel layer was hindering growth for LP in R_supp_ media, we tracked proliferation of LP in PVA inner layer gels, without the outer layer (**Figure 4**). We incubated the gels in R_supp_-10%, R_supp_-20% and MRS broth. Data obtained from the alamarBlue assay showed similar results to bilayer encapsulation for R_supp_-10% and R_supp_-20% (**Figure 4a, b**). MRS medium showed similar results as well, with a high decrease in fluorescence by day 2 (**Figure 4c**). From the brightfield images (**Figure 4d**), we confirmed that LP did not grow in the inner hydrogel layer cultured in R_supp_-10% and R_supp_-20%, whereas in MRS the bacteria formed colonies similarly to what was observed in bilayer hydrogels.

We next studied the growth of Cg in PVA gels. Prior to that, we confirmed that UV irradiation did not affect Cg growth in suspension (**Figure S5c**). Then we investigated Cg-PVA bilayer gels incubated in R-10% at 37°C using alamarBlue assay (**Figure S8a**). Bacteria did not grow at these conditions for up to 3 weeks. They also did not grow at 30°C in R_supp_-10%, R_supp_-20% and BHI broth (**Figure S8b-d**). We then compared the proliferation of Cg-PVA in hydrogels that only contained the inner layer at 30°C in BHI broth and R_supp_-20% (**Figure 5a, b**). We observed an increase in proliferation via AlamarBlue assay on day 1 for both BHI broth and R_supp_-20%. The brightfield images (**Figure 5c**) showed proliferation of Cg in the inner layer gels in R_supp_-20% with colony formation, which was not observed in BHI broth at the same timepoints.

**Figure 5.**
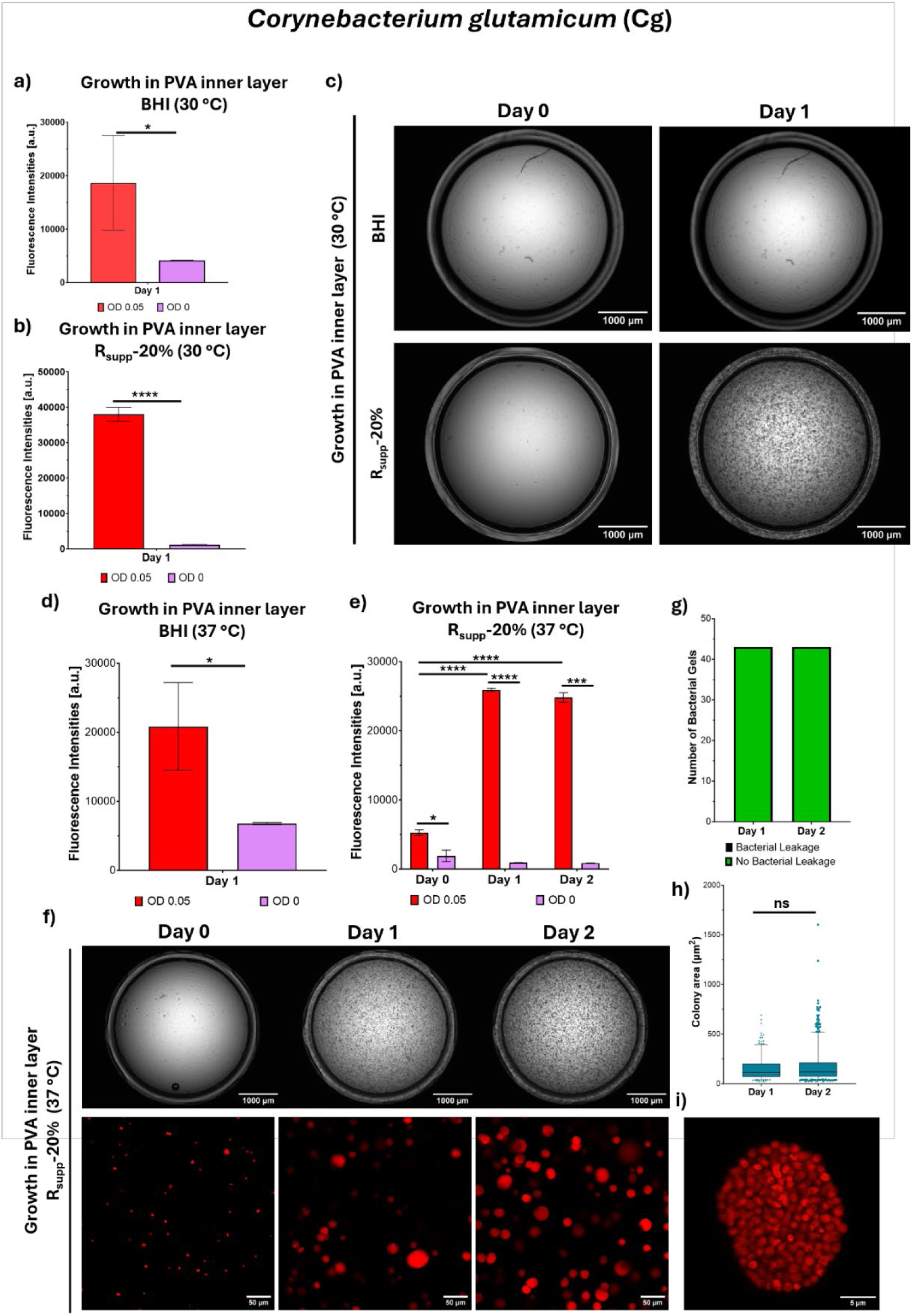
Cg encapsulated in PVA inner layer hydrogels grow in R_supp_-20% at 37°C. **(a)** Proliferation (manual gain 65, alamarBlue assay) of Cg-PVA inner layer gels in BHI broth at 30°C, **(b)** Proliferation of (manual gain 65, alamarBlue assay) Cg-PVA inner layer gels in R_supp_-20% at 30°C (mean ± SD). Differences among groups are indicated as follows: p-values <0.05 (*), p-values <0.01 (**), p-values <0.005 (***), p-values <0.001 (****). **(c)** Representative bright field images of Cg-PVA inner layer gels (scale bar: 1000 µm). **(d)** Proliferation of (manual gain 65, alamarBlue assay) Cg-PVA inner layer gels in BHI broth at 37°C, **(e)** Proliferation of (manual gain 65, alamarBlue assay) Cg-PVA inner layer gels in R_supp_-20% at 37°C, **(f)** Representative bright field images and Z-stack images (Z = 20 µm, Sum Intensities) of Cg-PVA inner layer gels (scale bars: 1000 µm and 50 µm, respectively). **(g)** Quantification of leakage from Cg-PVA inner layer gels in R_supp_-20% at 37° **(h)** Colony area quantification from the images (The line within the box signifies the median value, and the whiskers denote the range between the 5th and 95th percentiles, with outliers depicted as dots positioned above or below the whiskers). **(i)** Representative high magnification image of a single colony (Scale bar: 5 µm). Differences among groups are indicated as follows: p-values <0.05 (*), p-values <0.01 (**), p-values <0.005 (***), p-values <0.001 (****), ns = not significant.

Then, we studied the growth of Cg-PVA inner layer gels at 37 °C (**Figure 5d-i**). **Figure 5d** shows that Cg-PVA inner layer gels at 37°C incubated in BHI grew during the first 24 h. When incubating Cg-PVA inner layer gels in R_supp_-20% at 37°C, we observed rapid growth on day 1. This growth was maintained after day 2 (**Figure 5e**). From these constructs, we did not observe any leakage of bacteria up to day 2 (**Figure 5g**). Cg-PVA inner layer gels incubated in R_supp_-20% were also characterized via brightfield images and fluorescence microscopy (**Figure 5f-i**). On day 1, Cg were forming colonies and the number of colonies per field of view increased on day 2 (**Figure 5f**). Quantification of colony area showed that colonies formed on day 1 and 2 presented a similar average area (**Figure 5h**). High-magnification images showed that the production of mCherry was homogeneously distributed across the section of the colonies (**Figure 5i**).

For biocompatibility assessment, we performed studies with supernatants collected from Cg-PVA gels (**Figure 6a**) which we used in previous studies [7]. Cg-PVA gels were fabricated in inner layer hydrogels using the 96-well plate set up. First, we studied the viability of fibroblasts using LDH assay (**Figure 6b**). From this assay we quantified that only 10% of the cells had cell membrane disruption after addition of supernatants for 24 h. Alamarblue assay showed a lower reduction of the tetrazolium salt for cells that were in contact with bacterial supernatants (Cg-PVA) compared to cells in our controls (**Figure 6c**). Fluorescence intensities were similar to controls with pH adjusted to 5.5 (**Figure 6c**), which was the pH measured for the bacterial supernatants. Immunostaining revealed no appreciable differences in cell morphology (**Figure 6d, Figure S9**). We also carried out viability assays using monocytes via Live/Dead staining after addition of Cg-PVA supernatants (**Figure 6e, f, Figure S9**). Viability was similar for both bacterial supernatants and the empty hydrogel controls on day 1. Results showed that approximately 30% of cells remained viable on average, similar to the control condition with adjusted pH to 5.5 and significantly lower than the positive control.

**Figure 6.**
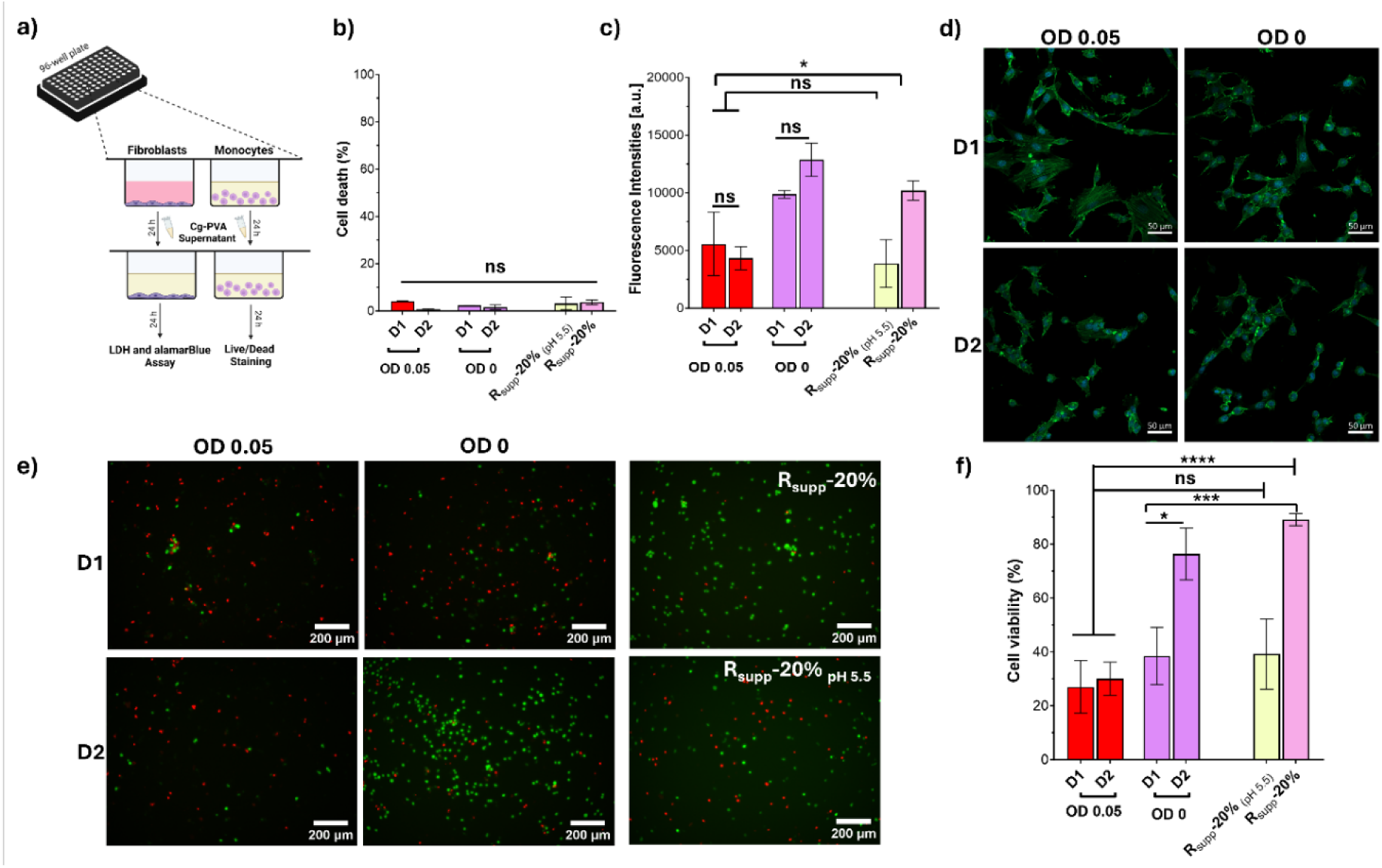
Biocompatibility studies on Cg-PVA hydrogels. **(a)** Representation of the procedures carried out to assess cytocompatibility of Cg-PVA supernatants. (**b**) Lactate dehydrogenase assay (LDH) on fibroblasts (NIH-3T3) with supernatants from day 1 and 2 of Cg-PVA gels OD 0.05 and OD 0. R_supp_-20% and R_supp_-20% with pH adjusted to pH 5.5 were used as controls, (**c**) fluorescence intensities measured during alamarBlue assay experiments on fibroblasts in contact with supernatants from day 1 and 2 of Cg-PVA gels OD 0.05 and OD 0. R_supp_-20% and R_supp_-20% with pH adjusted to pH 5.5 were used as controls. (d) Representative images of fibroblast morphologies after addition of supernatants from Cg-PVA gels OD 0.05 and OD 0 at days 1 and 2 (green: cytoskeleton, blue: nucleus, scale bar: 50 µm). (**e**) Representative images of live/dead staining of monocyte cells (mono-mac 6) in contact with supernatants of Cg-PVA gels OD 0.05 and OD 0 on days 1 and 2 where green depicts alive cells and red dead cells. Live/dead images from controls (R_supp_-20% and R_supp_-20% with pH adjusted to 5.5) are also shown. (**f**) Quantification of percentage of live cells was carried out from the images (mean ± SD, n ≥ 9, all statistically different comparisons are shown, all other comparisons are ns). Differences among groups are indicated as follows: p-values <0.05 (*), p-values <0.01 (**), p-values <0.005 (***), p-values <0.001 (****).

## 4. Discussion

Understanding how environmental and culture conditions affect ELMs and mammalian cells requires screening strategies coupling assays that track maintenance of ELMs while retaining compatibility with mammalian cells. To this end, we developed an ELM microarray in 96-well plate-based, to retrieve key information about ELM functionality together with the compatibility with the desired mammalian cells in an easy, robust way. The well-in-well architecture of the plates (μ-plate 96 well plate) used for ELM fabrication allowed us to assay bilayer and single layer formats. Further, we chose readouts that could be carried out in parallel using the 96-well plates and therefore, our screening methodology could track bacterial proliferation with alamarBlue and at the same time obtain brightfield images of the whole well and confocal images. Further, by using the supernatants we could measure the occurrence of bacterial leakage and test their compatibility with mammalian cells. From the images obtained, we were able to measure colony area as another parameter of bacterial growth. This methodology streamlines ELM proliferation investigation in different environmental conditions.

We started investigating EcN as it has been already reported to grow forming colonies in Pluronic-based ELMs when cultivated in RPMI medium [11]. Taking this into consideration, RPMI was chosen for initial experiments. To cope up with the nutritional requirements of mammalian cells, RPMI medium was supplemented with 10% FBS. Bacterial growth in R-10% **(Figure 3)** was noticeably lower compared to LB (control, **Figure S6**), which could be attributed to the higher amino acid to carbohydrate ratio present in RPMI as explained by Wijesinghe *et al.* [27]. These results were also in line with results from Rebai *et al.*, where growth curves of EcN in RPMI had overall lower OD600 compared to LB broth [28]. It is noteworthy that UV irradiation did not significantly affect EcN growth in R-10% or LB medium in suspension, as well as observed for LP and Cg (**Figure S5**). These findings were consistent with results reported by Li et al. for chemokine CXCL-12 secreting *Lactococcus lactis* [29]. Hydrogels should allow nutrient supply and gas exchange, so the encapsulated bacteria stay metabolically active and proliferating, as demonstrated by others [14, 16, 30, 31]. This was also the case for our ELMs as seen by an increase in fluorescence intensities of EcN-PVA bilayer gels in our proliferation studies (**Figure 3**). The colonies formed by EcN within our ELMs increased in size from day 1 to day 2 (**Figure 3**). These results were in agreement with the study from Liu *et al.* where EcN formed colonies in PVA-based magnetic ELMs [14] and from Riedel *et al.,* where EcN viability in PVA gels was maintained for up to 3 weeks [16]. Interestingly, the fluorescence intensities plateaued by day 2 and in contrast, the colonies grew from day 1 to day 2 (**Figure 3**). This can be explained by differences in metabolic activity of tightly packed colonies, nutrient competitive environment, and diffusion barriers. These stressed conditions could have induced microbial dormancy as discussed by Wood *et al.* [32, 33]. On the other hand, if bacteria had consumed all alamarBlue reagent available to them, then this could have also contributed to the observed plateau in alamarBlue assay. The nutrient content and its concentration in medium affect the colony sizes [34]. EcN colonies were considerably larger in R-10% medium compared to LB medium (**Figure 3 and S6**), which can be ascribed to its nutrient richness. In a similar study, *Phascolarctobacterium* grown with and without succinate on agar plates resulted in macro and micro colonies, respectively [35]. Nutrient and colony size correlations have been also observed in *Escherichia Coli*, forming smaller colonies when increasing glucose concentration in M63 medium [36]. In addition, Chacón *et al.* reported that *Salmonella enterica* colony sizes increase when changing the energy source to LB from acetate [37].

After successfully growing EcN-PVA in R-10% medium, we tried the same medium for LP-PVA bilayer gels with no success in terms of growth. The lack of growth in LP can be attributed to its 10-fold lower glucose concentration (2.0 g/L) compared to MRS broth. Interestingly, LP grew in R_supp_-10% (R-10% medium supplemented with extra glucose, HEPES buffer and sodium pyruvate) in suspension (**Figure 1**). However, LP-PVA bilayer gels did not proliferate for 14 days in the same medium (**Figure S7**). The alamarBlue assay has been previously reported to determine metabolic activity of LP in liquid culture [38] and also in confinement [39]. However, we had some discrepancies in our results. In agreement with Jun et al., the color of alamarBlue reagent and MRS broth mixture changed from blue to pale yellow. This can be explained by the acidic environment caused by the lactic acid produced by LP, which in turn, reacts with alamarBlue reagent [40]. To increase medium diffusion, we decided to eliminate the outer protecting layer. Additionally, we cultivated LP-PVA inner layer gels in a richer medium (R_supp_-20%). However, that did not increase bacterial growth, suggesting that LP might have other nutritional requirements. These observations show how our simple *in vitro* setup was able to identify the different requirements imposed by the chosen material, PVA-VS, which could not be overcome by using the physiologically-like conditions offered by the cell culture media. This suggests that further optimization of the material composition is required, or other nutritional requirements need to be included in the cell culture medium to achieve compatible conditions for LP growth in the PVA hydrogels used in this study.

For the third bacterial strain, Cg, the optimal medium and temperature for growth is 30 °C and BHI medium, respectively [41]. We opted for R-10% medium due to its similar glucose concentration (2.0 g/L) to BHI medium. Further, as the planned application for these ELMs are at physiological conditions, we monitored their growth at 37 °C in suspension. We found that these bacteria had overall slower growth at 37°C compared to 30°C (**Figure 1**). We tried to grow Cg-PVA bilayer gels at 30°C and with supplemented media (R_supp_-10% and R_supp_-20%) but observed no bacterial growth (**Figure S8**). However, the lack of growth observed for Cg-PVA gels in BHI medium at 30°C suggested that the 3D confinement environment and the possible lower oxygen diffusion through the bilayer gel format might have hindered proliferation. For example, Shyamkumar *et al.* observed lower glutamic acid production by Cg encapsulated in alginate constructs when increasing alginate concentration [42]. Cg is a facultative anaerobe and presents preferential growth in aerobic conditions, whereas it halts the growth in oxygen depriving conditions, although it can continue to survive in the presence of glucose [43]. Further, as measured previously by us 10 wt.% PVA-VS (our outer layer) has an oxygen permeability of 50 Dk [7] compared to the 80 Dk of water [44]. Considering this, we decided to eliminate the outer protecting layer to aid the oxygen diffusion through the Cg-PVA gels. We observed Cg growth forming colonies in Cg-PVA gels at both 30°C and 37°C (**Figure 5**). Interestingly, no colonies formed for Cg-PVA gels cultivated in BHI by day 1, which again stressed the critical role played by the chosen growth medium on growth behavior in confinement.

*In vitro* biocompatibility was assessed for Cg-PVA inner layer gel supernatants on fibroblasts and monocytes. Fibroblasts exposed to the Cg-PVA supernatants showed a decrease in fluorescence measured in alamarBlue assay (**Figure 6c**), which differed from values obtained for fibroblasts exposed to supernatants from empty gels (OD 0) and negative control (R_supp_-20%). Under oxygen depriving conditions, these bacteria produce succinic acid along with other acids making the ELM supernatant acidic [45], which could explain why the fluorescence values obtained for fibroblasts in contact with Cg-PVA supernatants were similar to a control medium with pH adjusted to 5.5 (acidic control). Similar results were obtained for viability studies performed on monocytes after addition of Cg-PVA supernatants (**Figure 6e, f**). Likewise, while co-cultivating human mesenchymal stem cells (hMSCs) and ELMs incorporating engineered *Lactococcus lactis*, Hay et al. obtained low *in vitro* cell viability of hMSCs and attributed it to medium acidification due to accumulation of lactic acid [46]. This stresses the idea that the dynamic nature of ELMs requires *in vitro* platforms that can systematically assess different parameters with time. Our 96-well plate-based screening method can tackle many of these parameters with the limitation that it only allows for relatively small volumes (70 μL capacity) and therefore, glucose consumption, waste accumulation and acidification of media might occur at a higher pace. Possible solutions could be included, such as better buffering systems or a more continuous media exchange during culture, which should be addressed independently for each ELM.

## 5. Conclusions

In conclusion, this work presents a methodology to systematically screen ELM culture conditions to assess the effect of environmental settings on ELM-host interactions *in vitro*. This setup allowed the parallel investigation of ELM proliferation in 20+ different microenvironmental conditions. We were able to assess bacteria proliferation in ELMs with three different bacterial species (EcN, LP and Cg) across 6 different cell culture media. We screened ELMs in bilayer and inner layer formats and measured the probability of bacterial leakage from the ELMs. We tracked colony growth via imaging with both light and fluorescence microscopies and were able to obtain high-magnification images with single bacteria resolution. Shape descriptors of bacterial colonies such as colony area were also implemented. *Corynebacterium* has appeared as a suitable species for ELMs with application to the eye as it is one of the main species in the eye microbiome. We took supernatants from Cg-PVA inner layer gels that grew in R_supp_-20% and studied cell biocompatibility. We selected a stromal, adherent murine cell line (NIH-3T3) and a non-adherent, immune, human cell line (mono-mac 6), and investigated their compatibility with our ELMs. Using this setup, we were able to find suitable culture media for two out of three bacterial strains investigated and perform initial biocompatibility studies in co-culture with mammalian cells. All in all, our setup has the potential to streamline research of ELMs in the biomedical field, helping in their translation from bench to bedside.

## Acknowledgements

ST would like to thank the Deutsche Forschungsgemeinschaft for funding through the grant Safe-LM (DFG GZ: TR 2001/1-1) and the Pharmaceutical Research Alliance Saarland for funding this research work. Authors also acknowledge the Fluorescence Microscopy Facility at the INM – Leibniz Institute for New Materials for help and advice with microscopy, and Lara Teruel Enrico for providing vinyl sulfonated -poly (vinyl alcohol). Authors thank Varun Tadimarri, Sourik Dey and Florian Riedel for generously providing mCherry expressing bacterial strains of LP, EcN, and Cg that were used in this study. Authors would also like to acknowledge BioRender for making it simple to create scientific illustrations of this work.

