## Supplemental material for "A screening setup to streamline *in vitro* Engineered Living Material cultures with the host"

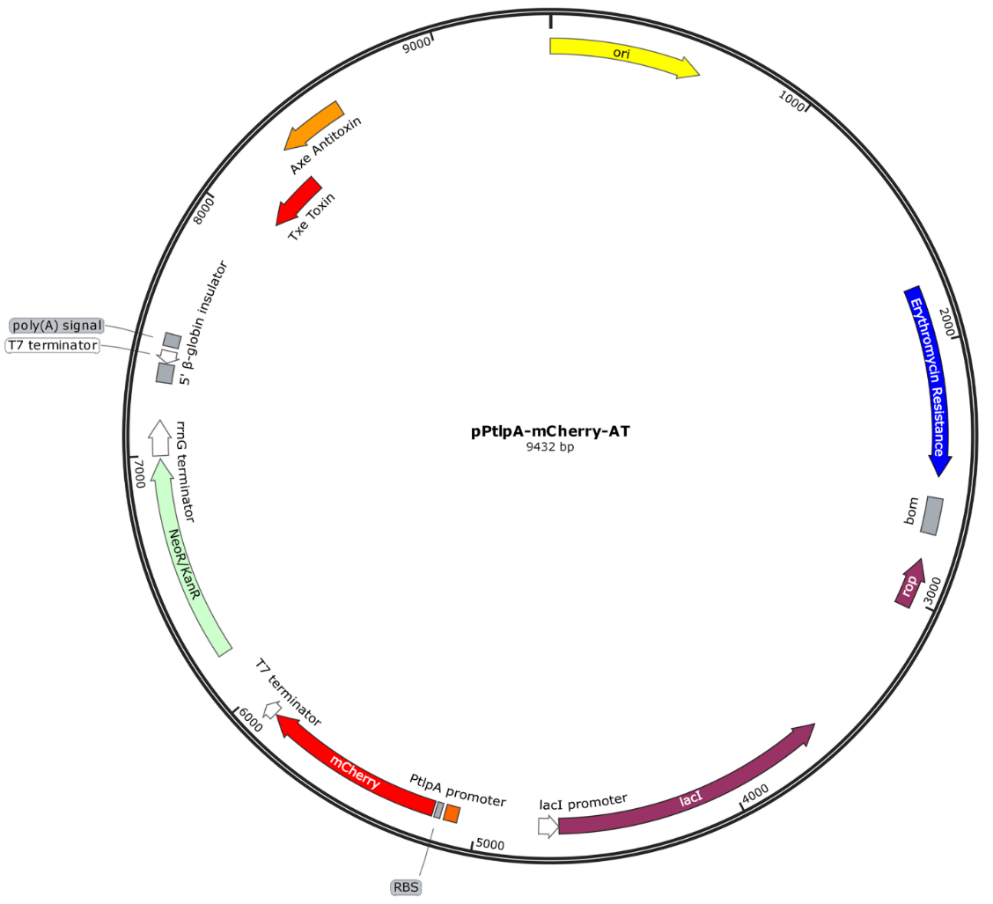

**Figure S1.** Plasmid map of the pPtlpA-mCherry-AT construct. The mCherry gene is constitutively driven by the PtlpA promoter in EcN 1917 and plasmid retention is facilitated by kanamycin supplementation.

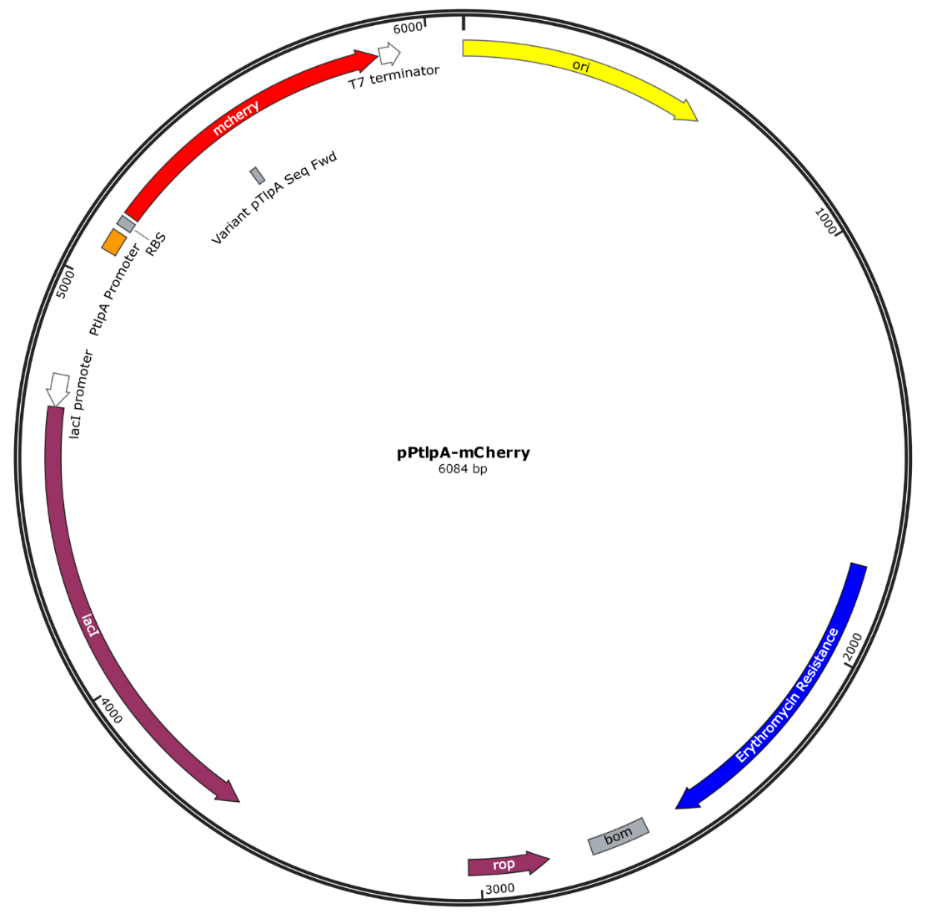

**Figure S2.** Plasmid map of the pTlp-mCherry construct. The mCherry gene is constitutively driven by the PtlpA promoter in LP WCFS1 and plasmid retention is facilitated by erythromycin supplementation.

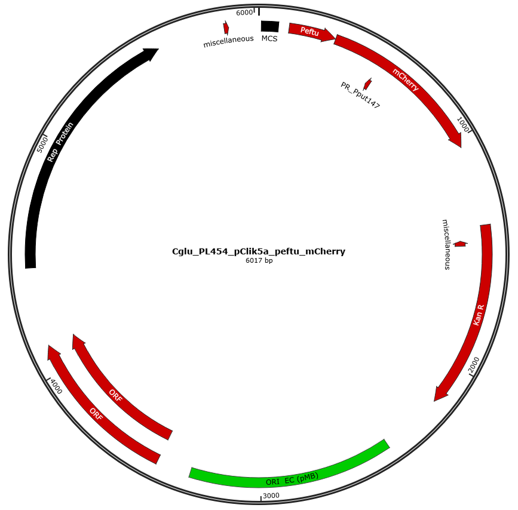

**Figure S3**. Plasmid map of the Cglu_PL454_pClik5a_peftu_mCherry construct. The mCherry gene is constitutively driven by the Peftu promoter in Cg and plasmid retention is facilitated by kanamycin supplementation.

**Table S1**. Individual volumes of inner layer components for EcN-PVA gels prepared in µ-plate 96 Well plates. Medium = R-10% or LB. All media were supplemented with kanamycin (50 µg/mL).

| EcN  Inner layer | Total volume | PVA-VS 10 % solution | PVA 10% solution | LAP in Medium | EcN suspension OD 0.5 in Medium |
| --- | --- | --- | --- | --- | --- |
| OD 0.05 | 300 μL | 142.5 μL | 7.5 μL | 120 μL (1.25 %) | 30 μL |
| OD 0 | 300 μL | 142.5 μL | 7.5 μL | 120 μL (1.25 %) | 30 μL (of only Medium) |

**Table S2**. Individual volumes of core components for LP-PVA gels prepared in µ-plate 96 Well plates. Medium = R_supp_-10% or Rs_upp_-20% or MRS Broth. All media were supplemented with erythromycin (10 µg/mL).

| LP Inner layer | Total volume | PVA-VS 10% solution | PVA 10% solution | LAP in Medium | LP suspension OD 0.5 in Medium |
| --- | --- | --- | --- | --- | --- |
| OD 0.05 | 300 μL | 142.5 μL | 7.5 μL | 120 μL (1.25 %) | 30 μL |
| OD 0 | 300 μL | 142.5 μL | 7.5 μL | 120 μL (1.25 %) | 30 μL (of only Medium) |

**Table S3**. Individual volumes of core components for Cg-PVA gels prepared in µ-plate 96 Well plates. Medium = R-10%, R_supp_-10%, Rs_upp_-20% or BHI. All media were supplemented with kanamycin (50 µg/mL).

| Cg Inner layer | Total volume | PVA-VS 10 % solution | PVA 10% solution | LAP in Medium | Cg suspension OD 0.5 in Medium |
| --- | --- | --- | --- | --- | --- |
| OD 0.05 | 300 μL | 142.5 μL | 7.5 μL | 120 μL (1.25 %) | 30 μL |
| OD 0 | 300 μL | 142.5 μL | 7.5 μL | 120 μL (1.25 %) | 30 μL (of only Medium) |

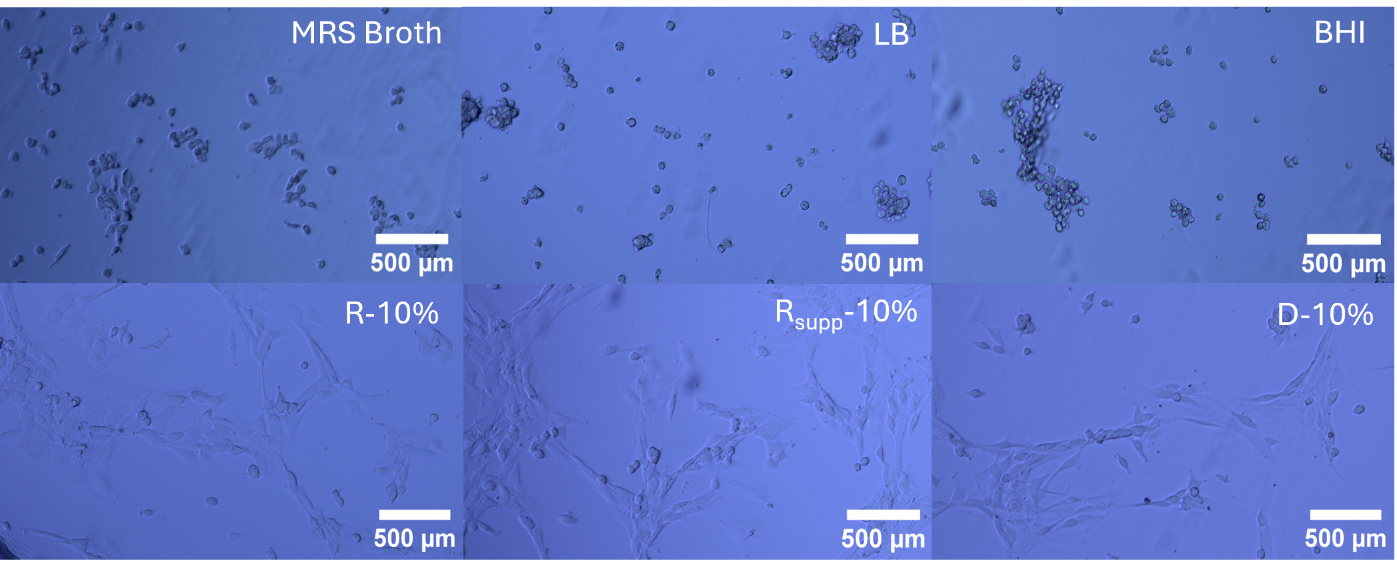

**Figure S4.** Representative Phase contrast images of fibroblasts in contact bacterial media. Phase contrast images from controls (R-10%, R_supp_-10% and D-10%) are also shown. Scale bar: 500 µm.

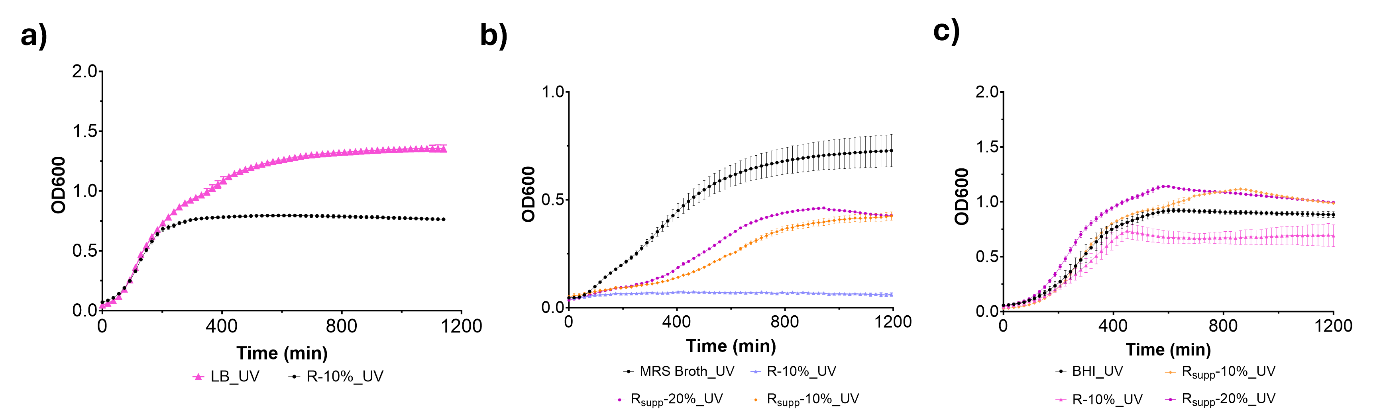

**Figure S5.** **UV irradiation does not affect growth of EcN, LP, and Cg in suspension.**

**(a)** Growth curves of UV irradiated EcN in suspension at 37°C in LB broth (pink line) and RPMI-10% FBS (R-10%, black line) **(b)** Growth curves of UV irradiated LP in suspension at 37°C in MRS broth (black line), R-10% (purple line), R_supp_-10% (orange line), R_supp_-20% (pink line) **(c)** Growth curves of UV irradiated Cg in suspension at 37°C in BHI broth (black line), R-10% (pink line), R_supp_-10% (light blue and yellow lines), R_supp_-20% (dark red line). For all plots data is represented as mean ± SD and n= 3 biological replicates.

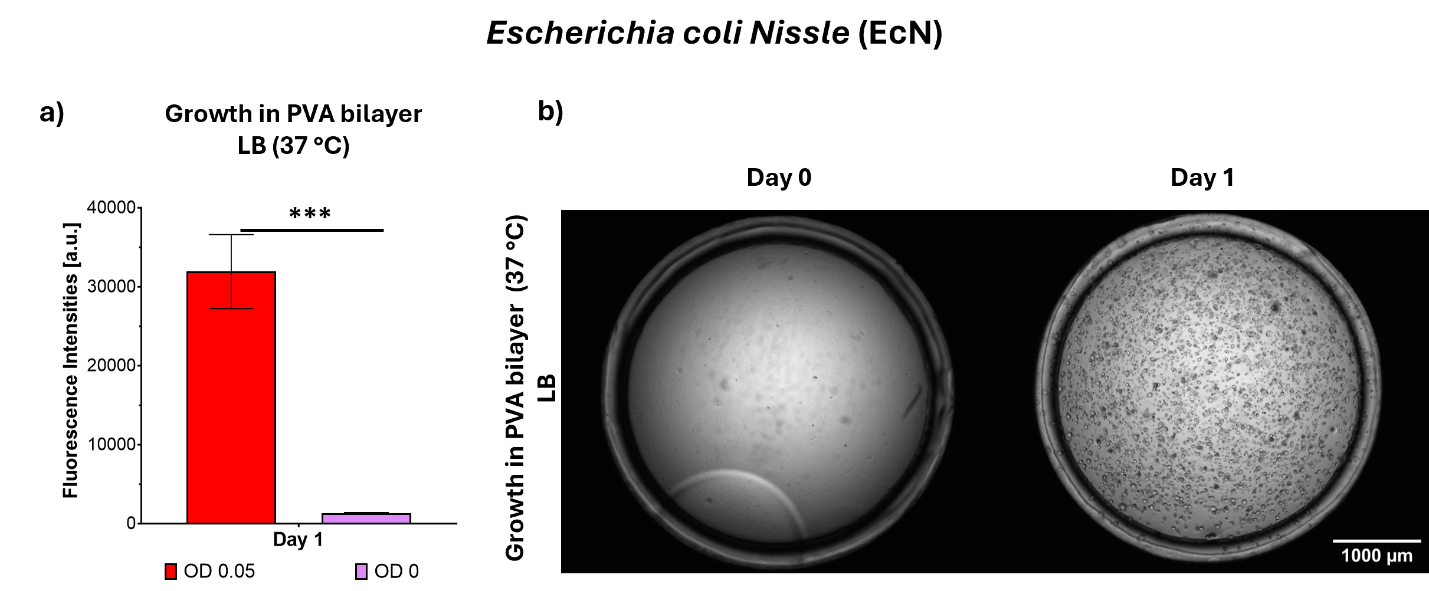

**Figure S6.** **(a)** Proliferation (manual gain 65, alamarBlue assay) of EcN-PVA bilayer gels in LB. **(b)** Bright field images of gels encapsulating EcN mCherry on day 0 and day, 1 respectively. Scale bar: 1000 µm.

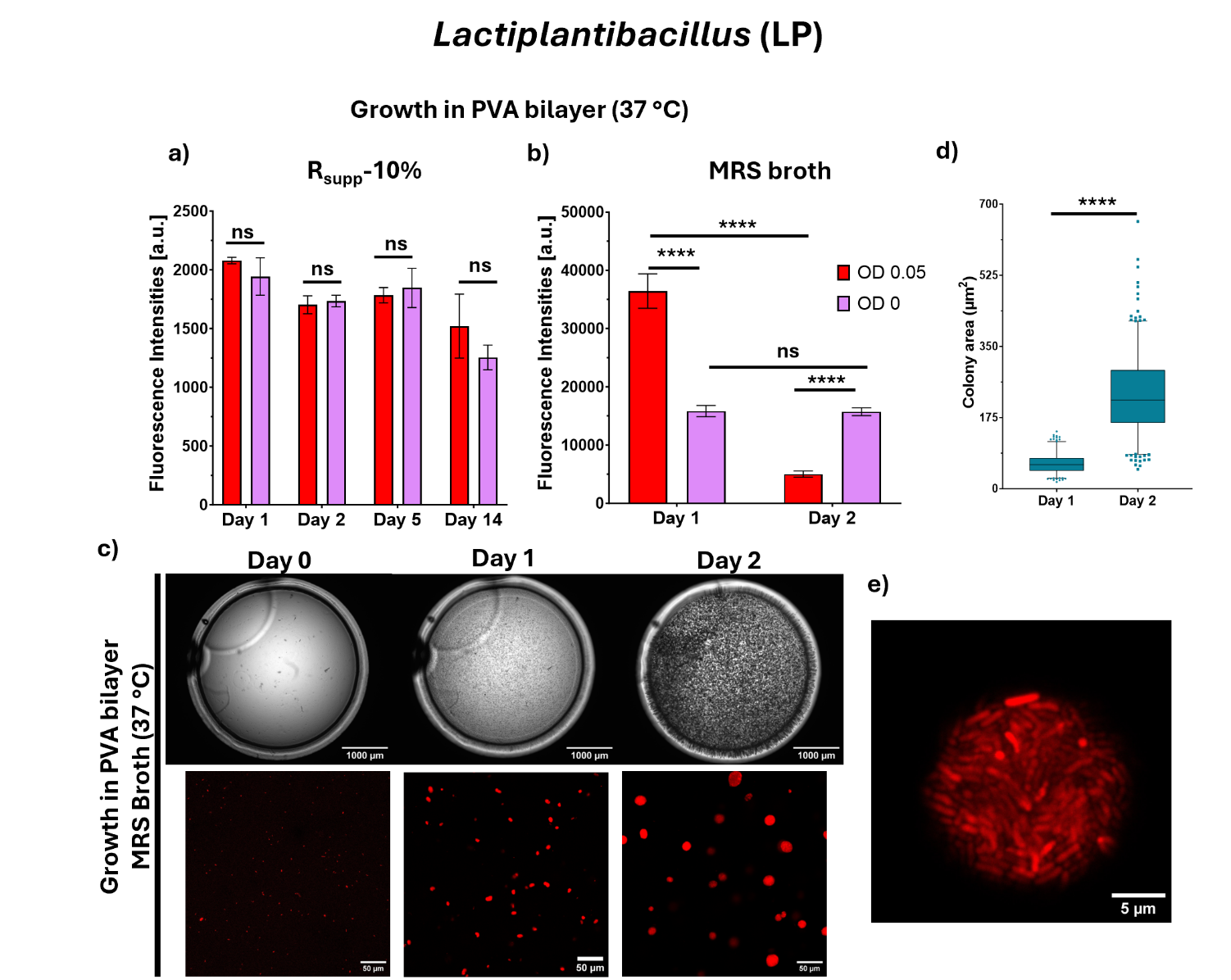

**Figure S7.** Proliferation (manual gain 65, alamarBlue assay) of LP-PVA gels in **(a)** R_supp_-10% and **(b)** MRS broth (n ≥ 3 biological replicates, mean ± SD). **(c)** Representative bright field images and Z-stack images (Z = 20 µm, Sum Intensities) of LP-PVA gels (scale bars: 1000 µm and 50 µm, respectively). **(d)** Colony area quantification from the images (The line within the box signifies the median value, and the whiskers denote the range between the 5th and 95th percentiles, with outliers depicted as dots positioned above or below the whiskers, n>200 colonies **(e)** Representative high magnification image of a single colony (Scale bar: 5 µm). Differences among groups are indicated as follows: p-values <0.05 (*), p-values <0.01 (**), p-values <0.005 (***), p-values <0.001 (****), and ns = not significant.

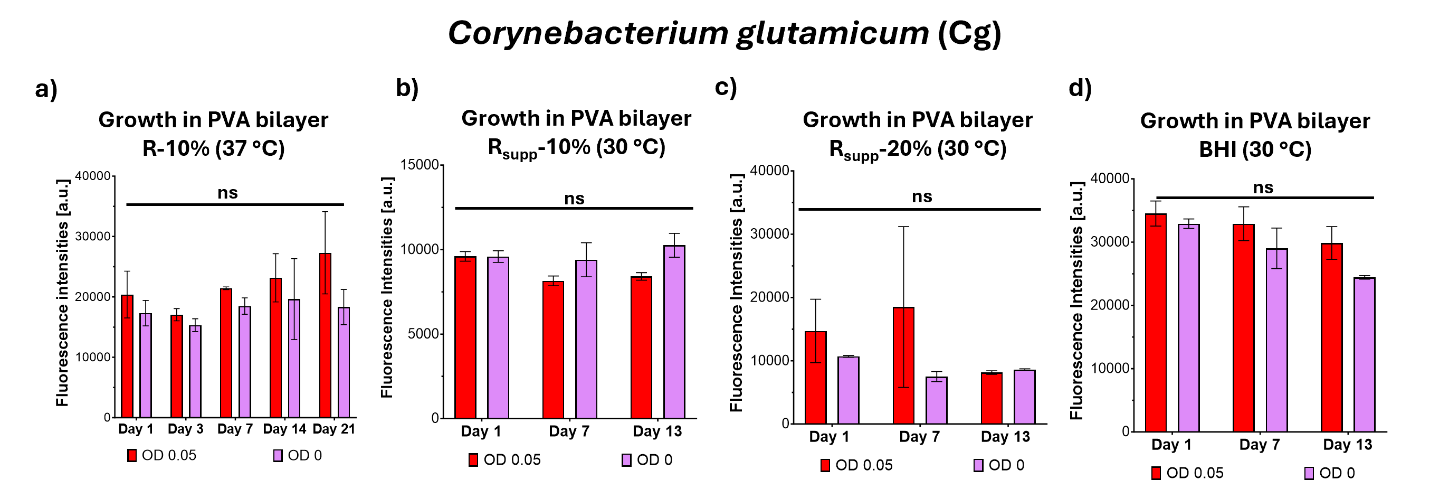

**Figure S8.** **(a)** Proliferation (manual gain 90, alamarBlue assay) of Cg-PVA bilayer gels in R-10% at 37°C, **(b)** Proliferation (manual gain 90, alamarBlue assay) of Cg-PVA bilayer gels in R_supp_-10% at 30°C, **(c)** Proliferation of (manual gain 90, alamarBlue assay) Cg-PVA bilayer gels in R_supp_-20% at 30°C, _supp_, **(d)** Proliferation of Cg-PVA bilayer gels in BHI broth at 30°C, Differences among groups are indicated as follows: p-values <0.05 (*), p-values <0.01 (**), p-values <0.005 (***), p-values <0.001 (****), and ns = not significant.

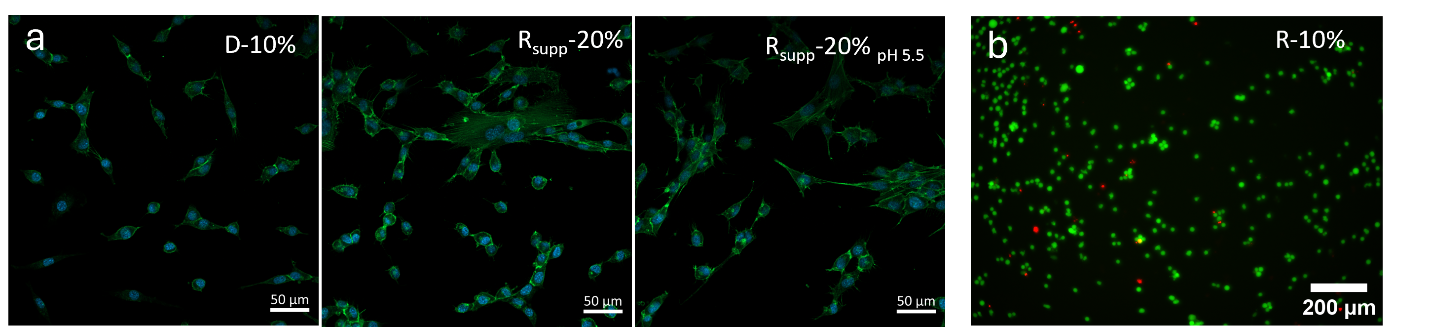

**Figure S9.** (**a**) Representative images of fibroblast morphologies in contact with controls (D-10%, R_supp_-20%, R_supp_-20% with pH adjusted to pH 5.5). (green: cytoskeleton, blue: nucleus, scale bar: 50 µm). (**b)** Representative images of live/dead staining of monocyte cells (mono-mac 6) in contact with R-10% where green depicts alive cells and red dead cells. Scale bar: 200 µm.
